## supplemental figures for "Expression of modified FcγRI enables myeloid cells to elicit robust tumor-specific cytotoxicity"

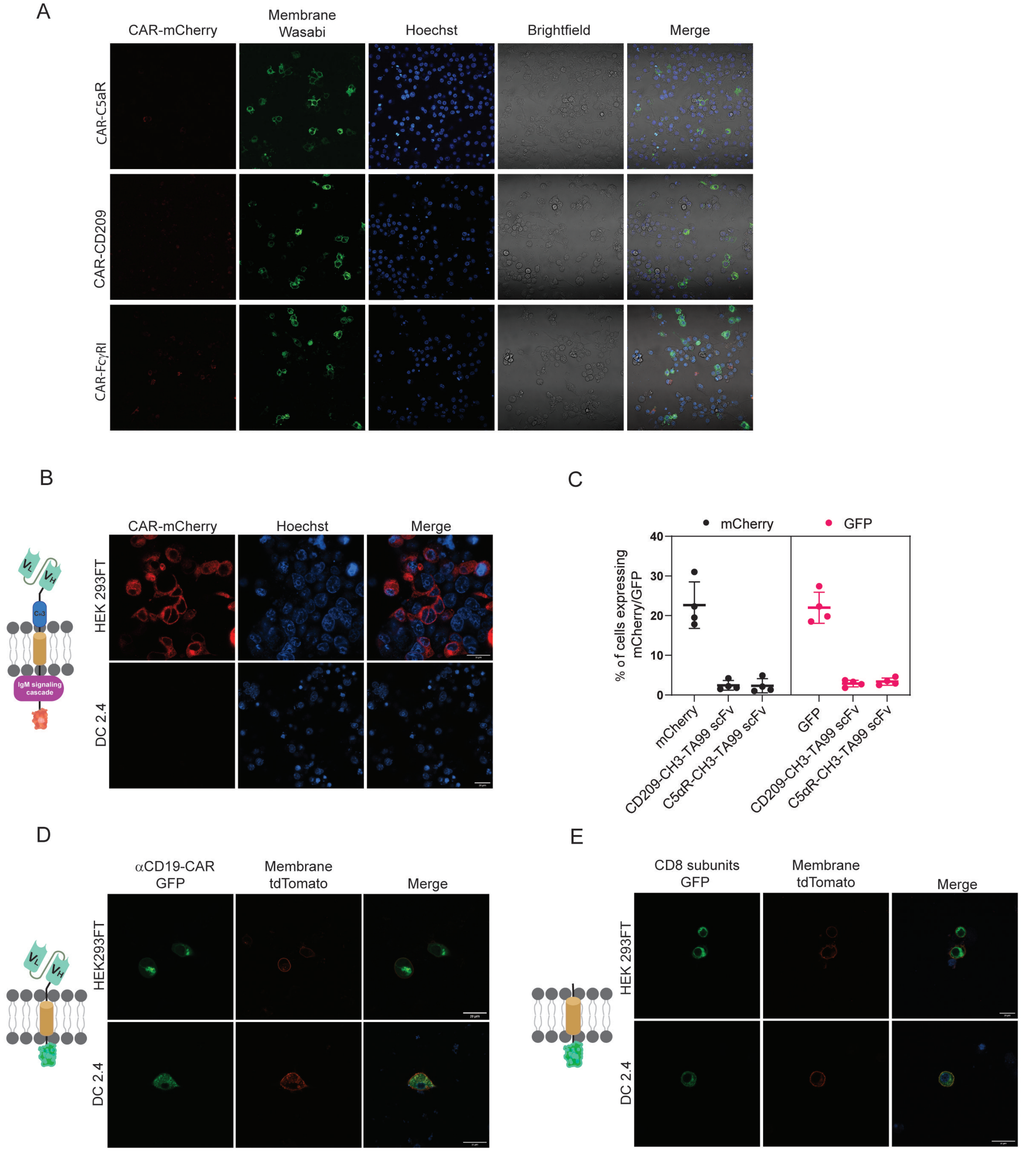


***Supplementary Figure S1: scFv is not expressed by myeloid cells.*** **(A)** Representative confocal images of RAW264.7 cells transfected with scFv-based chimeric receptors. **(B-C)** Representative confocal images **(B)** and mean expression percentages **(C)** of HEK293FT and DC2.4 cells 24 h following transfection with TA99 scFv. **(D-E)** Representative confocal image of HEK293FT and C2.4 cells 24 h following transfection with α CD19 scFv **(D)** and with CD8-transmembrane portion only **(E)**. Results are from one representative experiment out of at least three performed. Statistical significance was calculated using non-parametric t test

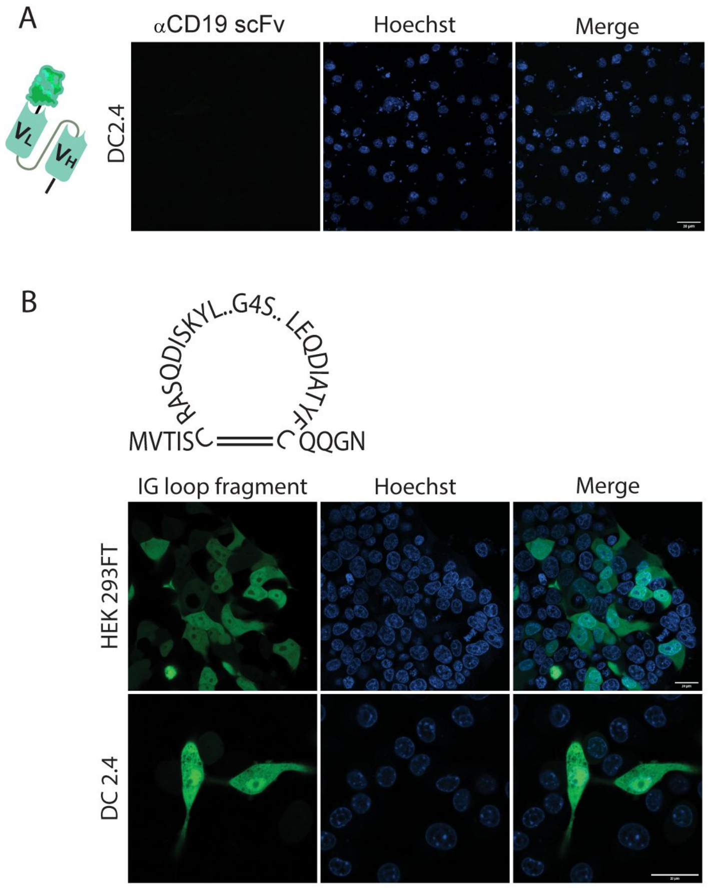


***Supplementary Figure S2. Immunoglobulin structure of scFv does not prevent degradation by myeloid cells.*** **(A)** Representative confocal images of DC2.4 transfected with α CD19 scFv fused to GFP at the carboxyl end[1]. **(B)** Upper: illustration of immunoglobulin loop fragment GFP. Lower: confocal microscopy images of HEK293FT and DC. Results are from one representative experiment out of at least three performed. [1]illustrations were created using BioRender.com

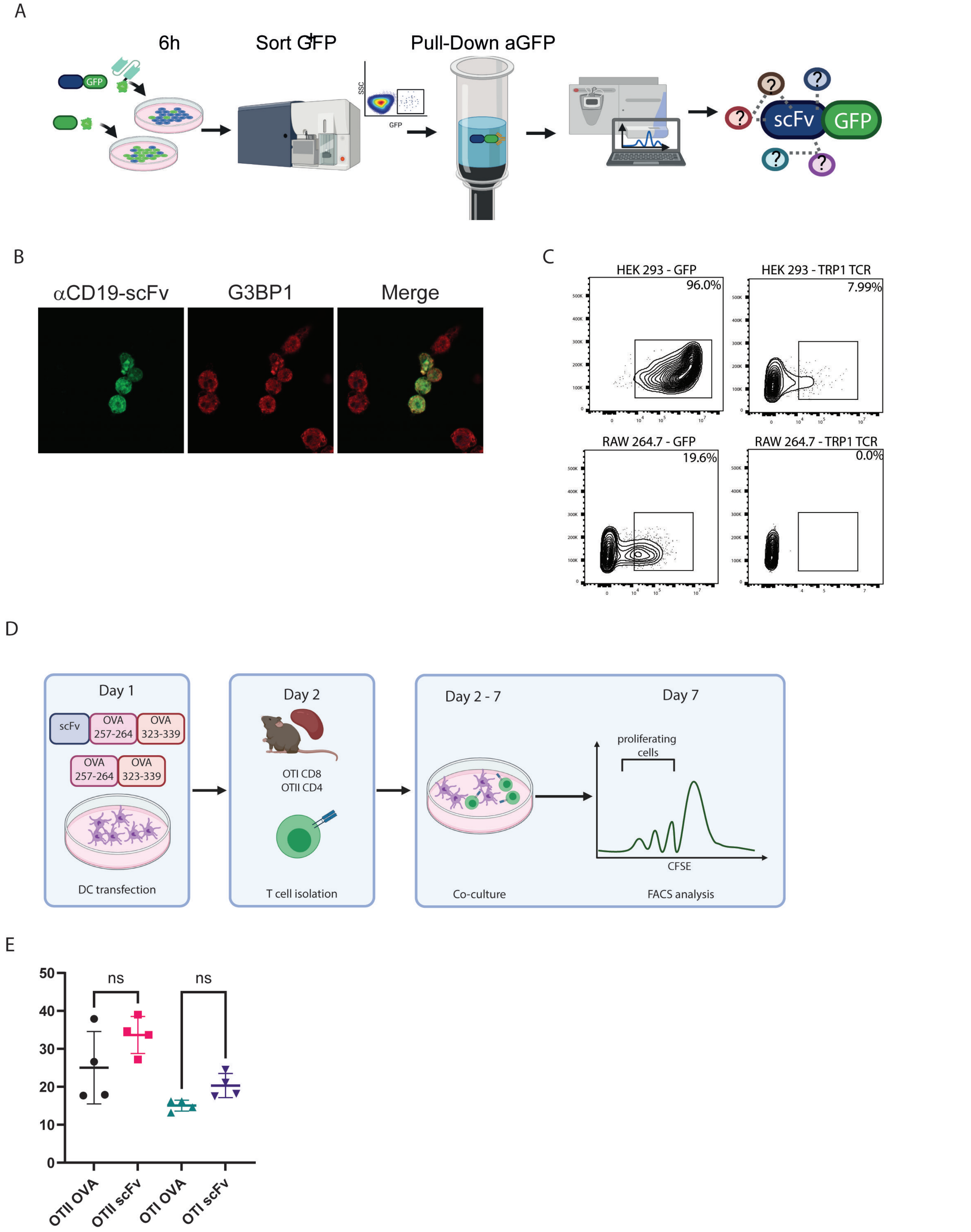


***Supplementary Figure S3. scFv induces ER stress response in myeloid cells.*** **(A)** Illustration of experimental design.[1] **(B)** Representative confocal staining for G3BP1 in DC2.4 cells 6 hours following transfection with αCD19 scFv. **(C)** Representative FACS analysis of HEK239FT and RAW 264.7 cells 24 hours post transfection with TRP1-TCR. **(D)** Illustration of experimental design.[1] **(E)** Representative analysis of CFSE dilution in CD8+ and CD4+ T cells following co-culture with ova conjugated scFv. Results are from one representative experiment out of at least three performed. Statistical significance was calculated using non-parametric t-test. [1]1Illustrations were created using BioRender.com

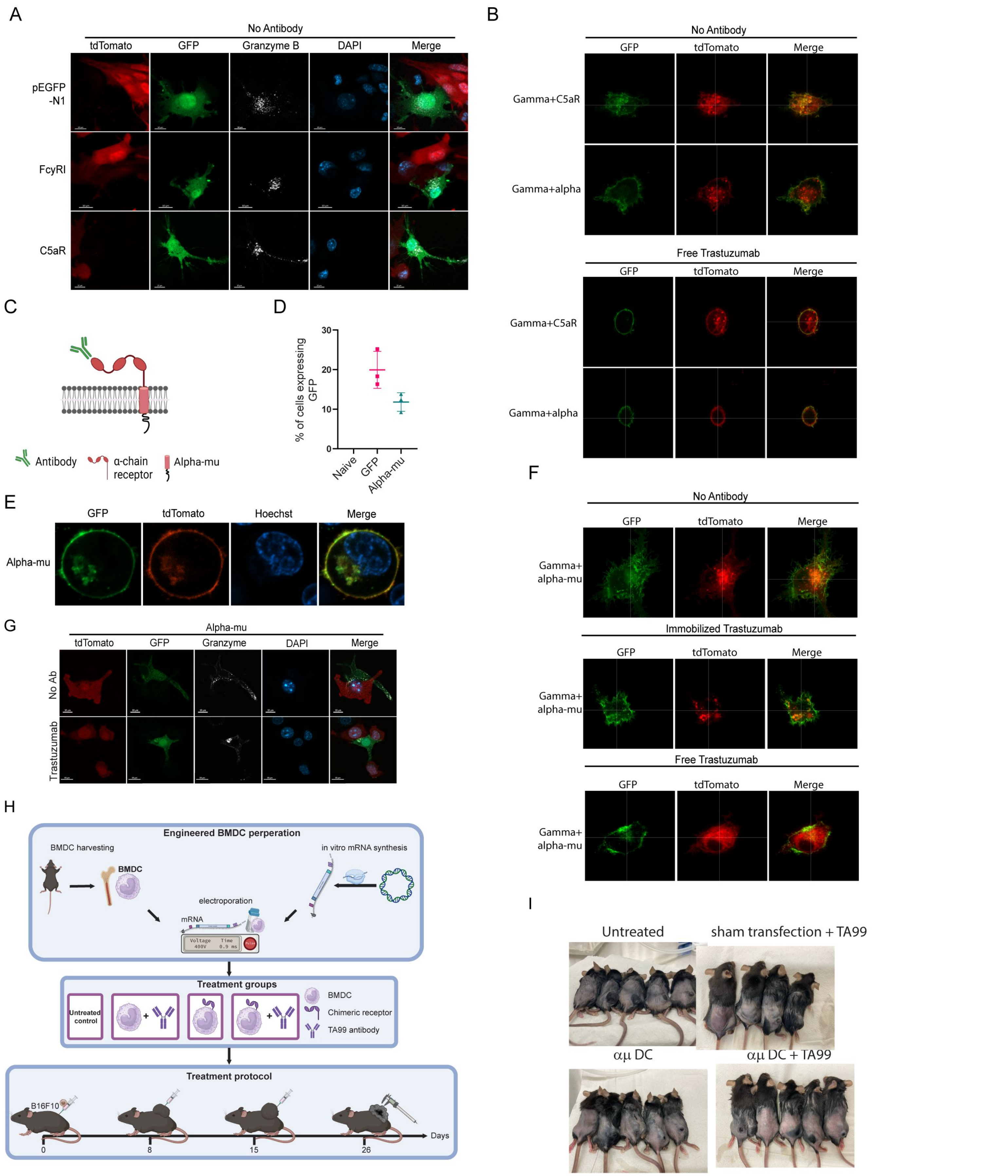


***Figure S4. FcγRI can be used as a scaffold to transmit IgM -induced signaling.*** **(A)** Representative confocal staining of Granzyme B in transduced RAW264.7 cells incubated overnight with 4T1 cells expressing human HER2 antigen. **(B)** Super resolution microscopy of RAW264.7 cells transfected with chimeric FcγRI molecules as such, or one hour after addition of free antibody. **(C)** Illustration of receptor design.[1]. **(D)** Mean percentages of RAW264.7 cells expressing chimeric receptor (n=4). **(E)** Representative microscopy of RAW264.7 cells co-transfected with GFP-fused chimeric receptor and mCherry-membrane protein. **(F)** Super resolution microscopy of RAW264.7 cells transfected with chimeric FcγRI molecules as such, or one hour after addition of free antibodies. **(G)** Confocal microscopy staining of GrB in RAW 264.7 cells co-cultured overnight with 4T1 cells expressing human HER2+. **(H)** Illustration of experimental setting.[1] **(I)** Representative photomicrographs of tumor-bearing mice 26 days after tumor injection. Results are from one representative experiment out of at least three independent experiments performed. [1]Illustrations were created using BioRender.com
